# Extinction times in diffusive public good population dynamics

**DOI:** 10.1101/424580

**Authors:** Gregory J. Kimmel, Philip Gerlee, Philipp M. Altrock

## Abstract

The co-evolutionary dynamics of competing populations can be strongly affected by frequency-dependent selection and population structure in space. As co-evolving populations grow into a spatial domain, their initial spatial arrangement, as well as their growth rate differences determine the dynamics. Here, we are interested in the dynamics of producers and free-rider co-evolution in the context of an ecological public good that is produced by a sub-population but evokes growth benefits to all individuals. We consider the spatial growth dynamics in one, two and three dimensions by modeling producer cell, free-rider cell and public good densities in space, driven by birth, death and diffusion. Typically, one population goes extinct. We find that uncorrelated initial spatial structures do not influence the time to extinction in comparison to the well-mixed system. We derive a slow manifold solution in order to estimate the time to extinction of either free-riders or producers. For invading populations, i.e. for populations that are initially highly segregated, we observe a traveling wave, whose speed can be calculated to improve the extinction time estimate by a simple superposition of the two times. Our results show that local effects of spatial dynamics evolve independently of the dynamics of the mean populations. Our considerations provide quantitative predictions for the transient dynamics of cooperative traits under pressure of extinction, and a potential experiment to derive elusive details of the fitness function of an ecological public goods game through extinction time observations.

**Author Summary:** Ecological public goods (PG) relationships emerge in growing cellular populations, for example between bacteria and cancer cells. We study the eco-evolutionary dynamics of a PG in populations that grow in space. In our model, public good-producer cells and free-rider cells can grow according to their own birth and death rates. Co-evolution occurs due to public good-driven surplus in the intrinsic growth rates and a cost to producers. A net growth rate benefit to free-riders leads to the well-known tragedy of the commons in which producers go extinct. What is often omitted from discussions is the time scale on which this extinction can occur, especially in spatial populations. We derive analytical estimates of the time to extinction in different spatial settings, and identify spatial scenarios in which extinction takes long enough such that the tragedy of the commons never occurs within the lifetime of the populations. Using numerical simulations we analyze the deviations from analytical predictions. Our results have direct implications for inferring ecological public good game properties from *in vitro* and *in vivo* experimental observations.

## Introduction

Heterogeneity and spatial patterns appear spontaneously in nature, on a wide range of spatial and temporal scales [1, 2, 3, 4]. Species are not randomly dispersed, but aggregate according to climate, predators, and resource levels, all of which can vary in both space and time. Structures of this type are, however, not always the result of external factors, but can also arise due to interactions between individuals of a population [5]. Thus, growing cell populations can be simultaneously driven by frequency-dependent and density-dependent selection [6], and the combination of the two mechanisms can lead to novel phenomena [7, 8]. Interactions between individual organisms are often mediated by their phenotypes, or strategies. In terms of reproductive success, the fitness of a certain strategy often depends on the frequency of other strategies present in the population. This frequency-dependence sets the stage for a game theoretic explanations of population dynamics. The ecological public goods game [3] describes a scenario in which a subpopulation releases costly factors, such as enzymes, into the environment, where they benefit both the producers, and non-producers.

The public goods game is played between producers of the public good and free-riders. Individuals either produce public good and thus ‘cooperate’, or only reap the benefits, i.e. free-ride and thus ‘defect’ [9]. This game has been studied by considering a group of *N* players [10], in which producers contribute the good at cost *κ* > 0 to itself. In the case that the benefit of the good is outweighed by the cost of production, free-riders will invade and outcompete the producers, which leads to the tragedy of the commons in which the overall population fitness declines as free-riders take over. This social dilemma-setting may also be observed in cancer cell growth kinetics [11], in which a subset of the population produces a growth factor (e.g. testosterone in prostate, EGF in lung cancer and PDGF in glioma [12]), while non-producing cells receive a potential benefit. This situation calls for explicitly modeling space, since the growth factor tends to be localized to producer cells and is transported by means of diffusion, which has a limited range.

Traditional chemotherapy protocols advise the maximum tolerable dose (MTD), which targets to eliminate drug-sensitive cells and often selects for drug-resistance. Relapse is then caused by proliferation of drug-resistant cells. As an alternative, adaptive therapy (AT) has recently been used successfully in cancer treatments [13]. It utilizes a variable dosing schedule to control (in theory) the diversity of the cancer. Proliferation of drug-sensitive cells allows for greater competition (e.g. contact inhibition, resource allocations) between cell types–this inhibits the proliferation of drug-resistant tumor cells. Clinical trials in breast [14], ovarian [15, 16] and prostate [17] have demonstrated that evolution-based AT strategies can be successfully employed, sometimes indefinitely and are superior to the standard MTD. Zhang et. al. in [17] employed a three population Lotka-Volterra model which consisted of cells requiring exogenous androgen (T+), cells which can produce androgen (TP) and cells which do not require androgen (T-). They showed via mathematical modeling the utility of adaptive therapy to control tumor growth rather than hit with a maximum tolerated dose. In a pilot clinical trial the effectiveness was verified by a large improvement in median survival. The T+ and TP cells act as free-riders and producers, respectively. The additional cell type T- is relevant in the case of therapy as it the drug-resistance clone. In this work, we consider interactions of type described between the two species of T+ and TP cells.

It has been shown that not just the type of game, but also how one defines the spatial grid and the type of individuals that can determine the outcome [18]. While spatial public goods games are well-studied, their evolutionary dynamics in growing populations has only recently been investigated in a non-spatial setting [8]. The time to reach an equilibrium point, which we denote “fixation time”, or “extinction time” in the case of a monomorphic equilibrium point, may depend critically on differences in net growth rates. Cooperation between cell lines was studied under varying substrate concentrations and it was observed that segregation occurred more readily when substrate was limited [19]. These spatial pattern formations occurred as the population moved into the whole domain. Once the population approaches capacity, competition should take over and the dominant clone should fixate. The experiments however focused on the behavior of the initial front type and showed the variation was due to available substrate. The timing of events has not been studied in great detail before, partly because standard tools in evolutionary game theory–such as the replicator equation–can describe growing populations (but usually do not) [20], and thus can not capture differences in net growth rates that result from frequency dependent selection in a public goods game [21]. However, these time scales play an important role biologically, especially if the time to fixation is longer than the lifespan of the population of interest. Tumor growth is a typical example, where the total tumor burden might kill the patient before one cell type outcompetes the other.

In this paper, we the important step to extend the logistic model considered in [8] to allow for spatial variation. We analyze spatial heterogeneity in up to three dimensions and show that the fixation time can be influenced by spatial heterogeneity. In particular, non-random initial conditions can cause large increases in the extinction time. We assume a linear public good function, which leads to a reduction in possible equilibria [22, 21]. We are primarily concerned with how spatial effects in the evolutionary dynamics of the public goods game impact the time to an extinction event. To approximate extinction times in spatial systems, we calculate an estimate of the time to reach a slow manifold and compute the time spent on this manifold. Numerical simulations can be in good agreement with analytically estimated extinction times, except when structured domains prevail in space. Highly structured domains arise, for example, if producers and free-riders occupy mutually exclusive regions in space. Such segregation pattern resemble a traveling Fischer wave at the intersection between the two populations. The agreement obtained highlights that these are in fact ‘pulled’ waves as opposed to ‘pushed’ waves [23]. Thus, one can explore the time a traveling wave of free-riders needs to move across the entire domain. The such determined time scales of the eco-evolutionary public goods game dynamics could then effectively be used to infer the underlying fitness functions that drive an empirical observation.

## Methods

Our analysis in this manuscript focuses on spatial populations, and is based on cell type specific growth and death rates. We focus on two sub-populations: public good producers and free-riders, and we are interested in the question of how spatial variations in producer and free-rider densities affect the long-term behavior of their dynamics, in particular the time to reach a possible equilibrium configuration. The analysis of this system is not straightforward because, although producer cells bear a cost and are thus expected to go extinct, their local concentration and the resulting fluctuations in public good availability can influence the dynamics in interesting ways. Regardless of dimensionality, we show that any initial spatial variability is transient and equilibrates to a spatially homogenous solution. Therefore, a well-mixed ODE-model can be sufficient to analyze the long-term behavior of the spatial system. We construct a coupled dynamical system which models the behavior of public good producers and free-riders and the spatial distribution of public good (growth factor) in time and space. We derive slow manifold solutions which allow us to predict the time to reach an equilibrium (fixation or extinction time) for a wide range of parameters. Our calculations become particularly useful when extinction times are easily observable in experimental systems. Then, using multiple observations of producer cell extinction over time, one could infer the underlying parameters that drive overall fitness differences.

### Ecological public good dynamics in space

In the simplest setting we can assume that producer cells (*U*) and free-riders (*V*) are closely related cell types experiencing the same baseline growth rate *α* and the potentially different death rates *μ_U_, μ_V_*. Next, we assume that the tumor public good, produced by *U* cells, has a linear positive effect on the growth rates, in the form of an additive benefit-to-growth-rate proportional to the local public good or growth factor density *G*. This growth factor density is determined by the local producer cell density. *G* is produced by *U* cells at a cost to their growth rate κ, at a rate *ρ*, and it is consumed by *U* and *V* cells alike at a rate *δ*. The diffusion rate of the growth factor that acts as a dynamics ecological public good is Γ*_G_*.

We begin by considering the population game between producer and free-rider cells at respective densities *U*, and *V* in space and time. The cells are assumed to reside and grow on a spatial domain [0, *L*]*^n^* ⊂ ℝ *^n^*, where *n* = 1, 2, 3 is the dimension of the system. We assume that the domain has no-flux boundary conditions, that is cells cannot enter or leave the domain. We assume that growth (proliferation), death and competition processes are purely local and that migration (determined by the cell type specific diffusion coefficients Γ*_U,V_*) is isotropic and involved only with nearest neighbors. We then obtain the following set of coupled PDEs that model the distributions of producer and free-rider cells and the concentration of public good in time and space:

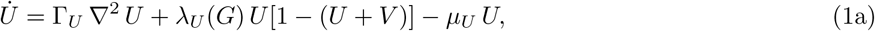

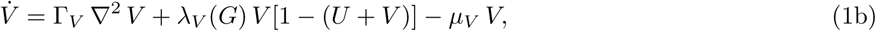

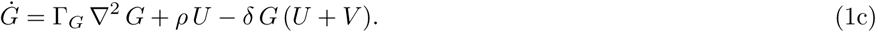

Here, the respective growth rates are

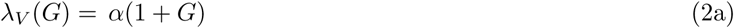

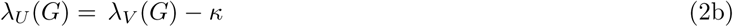

In absence of growth factor concentration, producer cells experience a growth rate detriment in amount of *κ*.

All important parameters and their baseline values are summarized in Table 1. The typical cell size is on the order of micrometers. Thus, in an attempt to simulate many cells, we focused on spatial domain ranges of *L* = 0.1–10 cm. The length of time for a cell cycle is potentially highly variable. A typical cell cycle could range from hours, to days, to weeks, and the proliferation (growth) rate is typically higher than the death rate [8].

**Table 1:** Dimensional parameters used in the model given by Eq. (1). The unit cc^-1^ means per cell cycle.

| Parameter | Symbol | Typical value |
| --- | --- | --- |
| Producer's diffusion coefficient | $\Gamma_U$ | $10^{-9} \text{ cm}^2/\text{s}$ |
| Free-rider's diffusion coefficient | $\Gamma_V$ | $10^{-9} \text{ cm}^2/\text{s}$ |
| Public good's diffusion coefficient | $\Gamma_G$ | $10^{-6} \text{ cm}^2/\text{s}$ |
| Cellular intrinsic growth rate | $\alpha$ | $1 \text{ cc}^{-1}$ |
| Producer's death rate | $\mu_U$ | $< 1 \text{ cc}^{-1}$ |
| Free-rider's death rate | $\mu_V$ | $< 1 \text{ cc}^{-1}$ |
| Public good production cost | $\kappa$ | $0.1 \text{ cc}^{-1}$ |
| Public good production rate | $\rho$ | $500 \text{ cc}^{-1}$ |
| Public good consumption rate | $\delta$ | $500 \text{ cc}^{-1}$ |
| Characteristic length of spatial domain | $L$ | $10 \text{ cm}$ |

We can construct the following non-dimensional form of the spatial model. In the original model formulation are eight parameters and three initial conditions *U*_0_(*x*), *V*_0_(*x*), and *G*_0_(*x*). With appropriate choices we can reduce the total number of relevant parameters to six dimensionless parameters. Although there are many choices for the set of dimensionless parameters; we here exploit that the time scale of the dynamics for *G* may be much faster than the time scales of the dynamics of *U* and *V* [8, 24]. After appropriate rescaling, we can use the dimensionless parameters of the non-dimensional system given in Table 2.

**Table 2:** Non-dimensional parameters used in the model given by Eq. (3). *ε*_exit_ is used to determine the extinction or fixation events.

| Dimensionless parameter | Symbol | Identity | Typical value |
| --- | --- | --- | --- |
| Producer's diffusion coefficient | $\gamma_U$ | $\frac{\Gamma_U \delta}{\Gamma_G \alpha}$ | 0.5 |
| Free-rider's diffusion coefficient | $\gamma_V$ | $\frac{\Gamma_V \delta}{\Gamma_G \alpha}$ | 0.5 |
| Producer (PG independent) birth rate | $a$ | $1 - \frac{\kappa}{\alpha}$ | 0.9 |
| Cell (PG dependent) birth rate | $b$ | $\frac{\rho}{\delta}$ | 1.0 |
| Producer death rate | $c$ | $\frac{\mu_U}{\alpha}$ | 0-1 |
| Ratio of free-rider to producer death rate | $r$ | $\frac{\mu_V}{\mu_U}$ | $> 0$ |
| Ratio of cell birth rate to consumption rate | $\epsilon$ | $\frac{\alpha}{\delta}$ | $2 \times 10^{-3}$ |
| Neighborhood of a fixed point | $\varepsilon_{\text{exit}}$ | | $10^{-8}$ |

We also introduce dimensionless time *τ* = *α t*. Space is scaled via 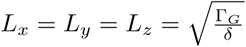 and *S* = *G δ/ρ*, which leads to non-dimensional domain lengths between 1 and 10^3^. In our notation, the “dot” then means differentiation with respect to dimensionless time *τ* (instead of *t*), and ∇ is the differential operator with respect to the rescaled spatial variables. Then we arrive at the dimensionless system

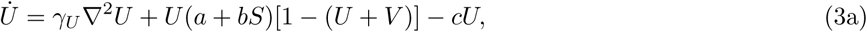

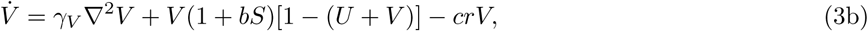

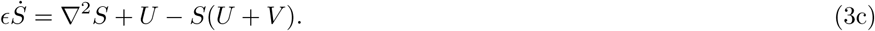

Turning to a dimensionless framework allows to obtain insights that are more readily apparent in this setting. For example, the public good consumption rate is typically much faster than the proliferation rate, *ϵ* ≪ 1, and we thus spatial equilibration of the public good *G* occurs relatively fast. Similarly, we can immediately see from the dimensionless system that the ratio of death rates between cell types, *r*, is an important quantity that determines the fate of cooperation, especially provided the ratio of death rate to proliferation rate in producer cells, *c*, is small.

## Results

### Spatial variation is transient regardless of dimensionality

What is the impact of variability in initial conditions? To address this question, we investigated the dynamics of the system (1) in one, two and three dimensions in its non-dimensional form (3). The non-dimensional length used was *L* = 10 for all spatial dimensions (*n* = 1, 2, 3). To solve numerically (3), we discretized the domain into grid points. The grid points were then given initial concentrations of the amount of producer, free-rider and public good present. The distance between grid points, or the spatial step size, was chosen to be no bigger than Δ*x* = 0.5. We tested smaller grid sizes, but found no significant changes in the dynamics, only in total CPU time. We solved the PDE using a Crank-Nicolson scheme with a time step size Δ*t* = 0.1 [25]. Unless specified differently, we set *r* = 1, i.e. we assumed that the two types had equal death rates. Then, 100 simulations were used to calculate summary statistics of the time to reach the neighborhood of a stable fixed point, with an exit criterion based on distance to the stable fixed point *d*(*U*, V\**) < *ε*_exit_, where the value *ε*_exit_ = 10^−8^ was used.

In all settings of different spatial dimension, we were interested in four types of initial conditions that define the initial cell density (amplitude) at every grid point: (1) Uniformly distributed values between 0 and 1, (2) domain wall (step function), (3) unimodal (e.g. a Gaussian perturbation), and (4) a bi-modal perturbation. Examples of the four initial condition types in 1D are shown in Figure 1. To examine the stability of the more structured density distributions (2)-(4), we also tested the impact of spatial noise by introducing a random deviation of the cell density in each point in space, which was chosen no greater than 10% of the max amplitude at each grid point.

**Figure 1:**
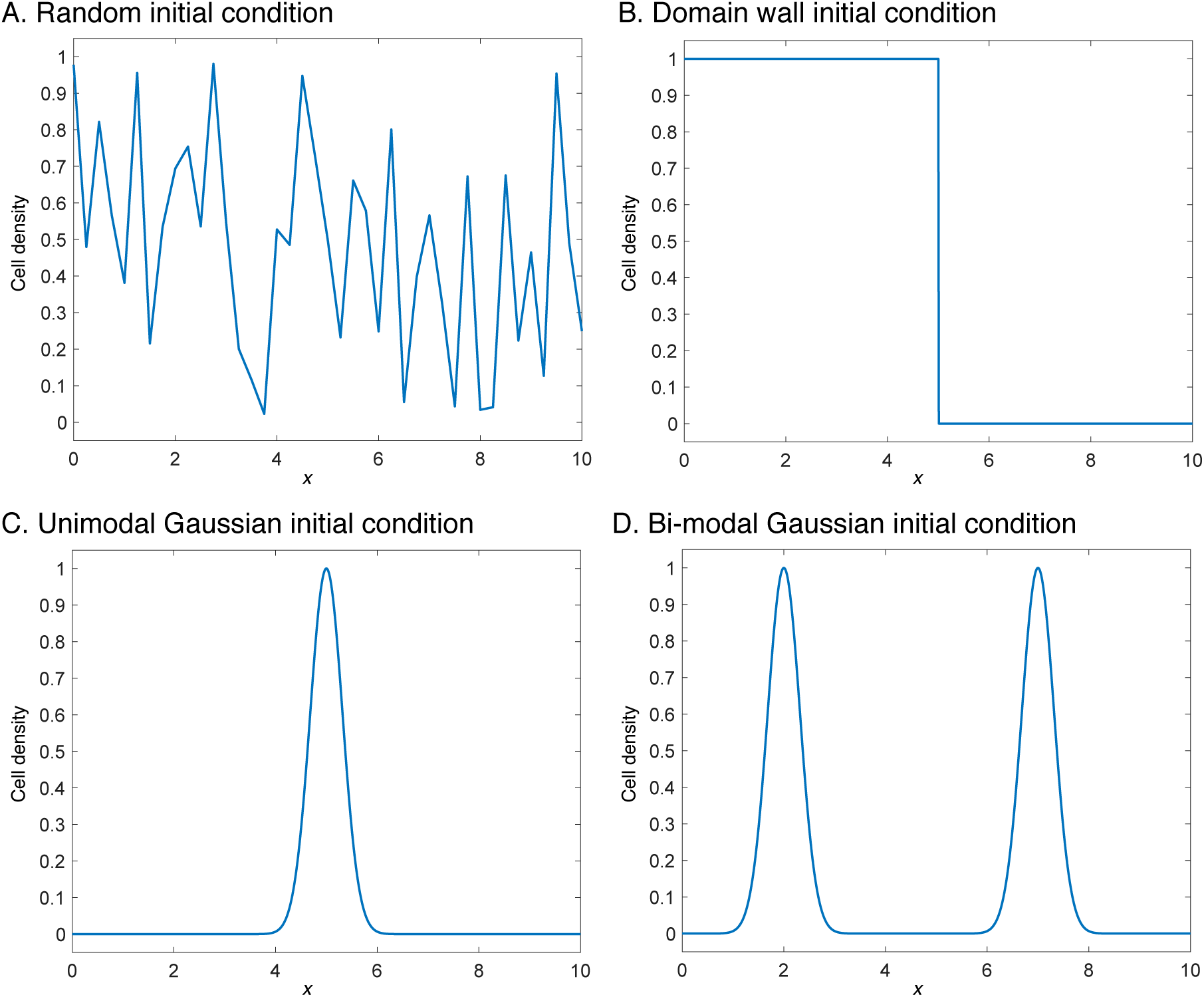
Initial conditions for cell density in one dimension. **A**: Values are assigned by drawing from a standard uniform distribution. **B**: Domain wall-type initial condition where the concentrations are separated. **C**: Unimodal condition using a Gaussian perturbation. **D**: Bi-modal initial condition using two separated Gaussian perturbations. In numerical simulations, all initial conditions were given a small random noise term.

Under the assumption of fast diffusion of cells into space, a spatial perturbation typically equilibrates along the spatial domain faster than an average cell cycle length. Figure 2 shows the temporal evolution of a typical simulation run, with a random initial condition being drawn from a standard uniform distribution on each grid point. The large oscillations in initial cell densities were damped during the first cell cycle. However, strong deviations from homogeneity persisted past *t* = 10 cell cycles. By the 100th cell cycle, the system essentially equilibrated and the remaining time was taken to reach the *ε*_exit_-radius (needed for the exit criterion). In this example, the exit condition was met at *τ* = 369.23. The average cell concentrations are shown in the second subplots and shows the phase diagram for the average cell concentrations. It is clear, that the average quickly reaches the slow manifold and spends most of its time traveling along it. The final snapshot shows the slow manifold, calculated from the ODE model in red. Though the model is clearly spatially dependent, the average cell population still rapidly approaches the slow manifold of the spatially averaged cell populations. It is clear that spatial variations can influence the time to extinction, however simulations have shown that spatial heterogeneity of final states do not exist.

**Figure 2:**
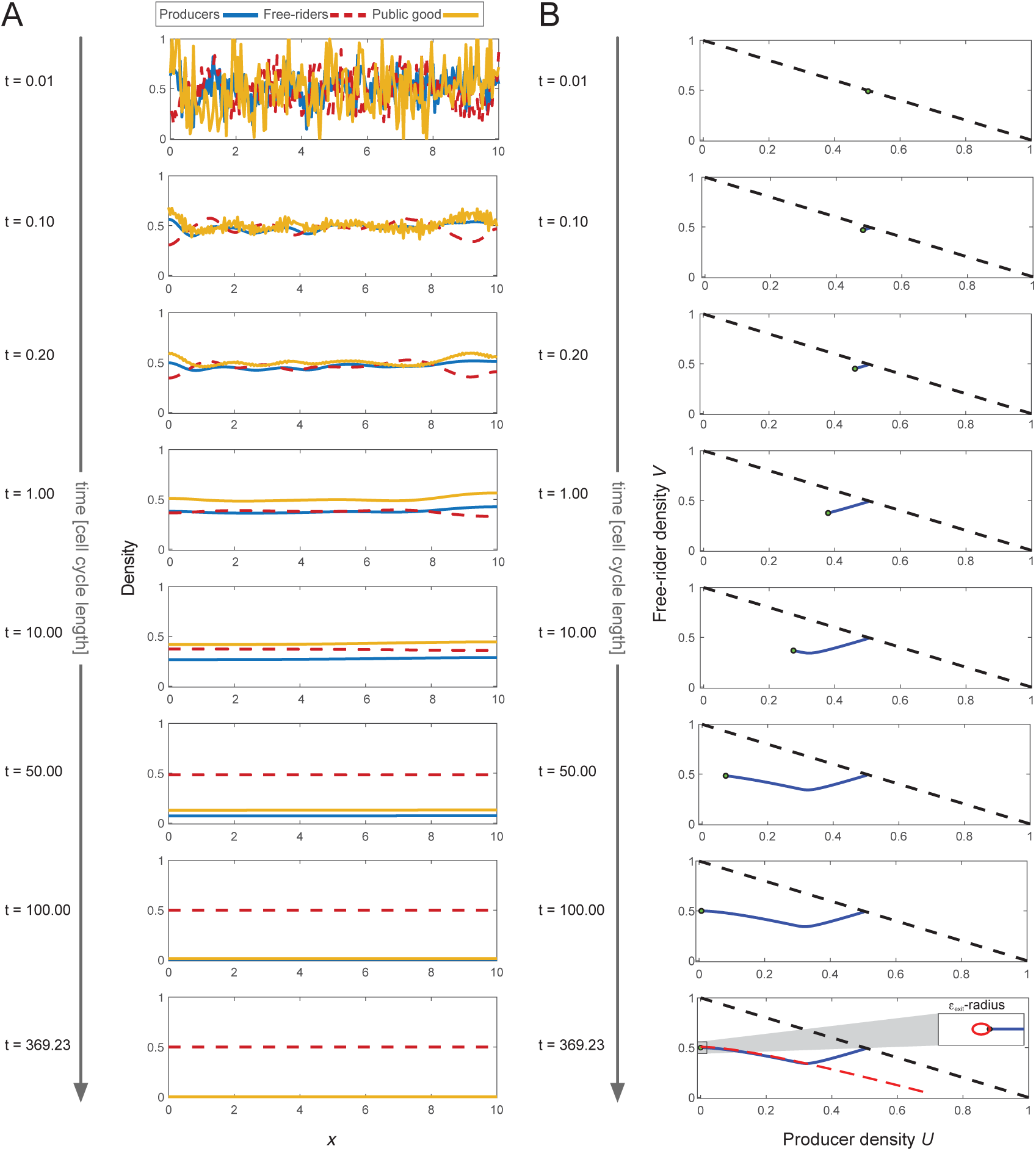
Typical 1D simulation leading to producer extinction. **A**: Snapshots of the system, represented by concentration of producer cells (*U*), free-riders (*V*) and growth factor (*G*) over time, measured in cell-cycle length (time advances top to bottom). The population game is played in 1D, the panels show the concentrations in space. **B**: Corresponding trajectory of producers and free-rider densities in their phase space (*U* = *V* on the black dashed line). Due to cell motility, the system reaches the slow manifold (red dashed line in bottom panel) fast, and spends most of the time traveling along the slow manifold. The bottom panel shows the *ε*_exit_ = 10^−8^-neighborhood exit criterion. Dimensionless parameters used: *c* = 0.5, and *γ_U_* = *γ_V_* = 0.5, *a* = 0.9, *b* = 1, *c* = 0.5, *r* = 1, *ϵ* = 2 × 10^−3^.

All numerical solutions approached spatially homogeneous solutions consisting of only free-riders within finite time, under our parameter assumptions, regardless of dimensionality. Using random initial conditions, we found that the average time to extinction was also independent of the dimensionality. However, comparing the numerical results shown in Figures 3 A, B, one can observe an apparent decrease in variance of the fixation (extinction) times as the dimensionality increases. This change in fixation time variation can be attributed to the increased number of grid points in higher dimensions within the same constant domain length (a *L* unit length line vs. a *L* × *L* unit square, etc.). We verified this attribution by running random initial conditions for all dimensions using a fixed total number of grid points (around 100 total points), regardless of dimensionality. If we compare the fixation time distributions of Figures 3 B and C, we can observe that the variance of the extinction time distributions become indistinguishable across dimensions once the number of spatial sites (grid points) with non-zero cell density is kept constant in all dimensions.

**Figure 3:**
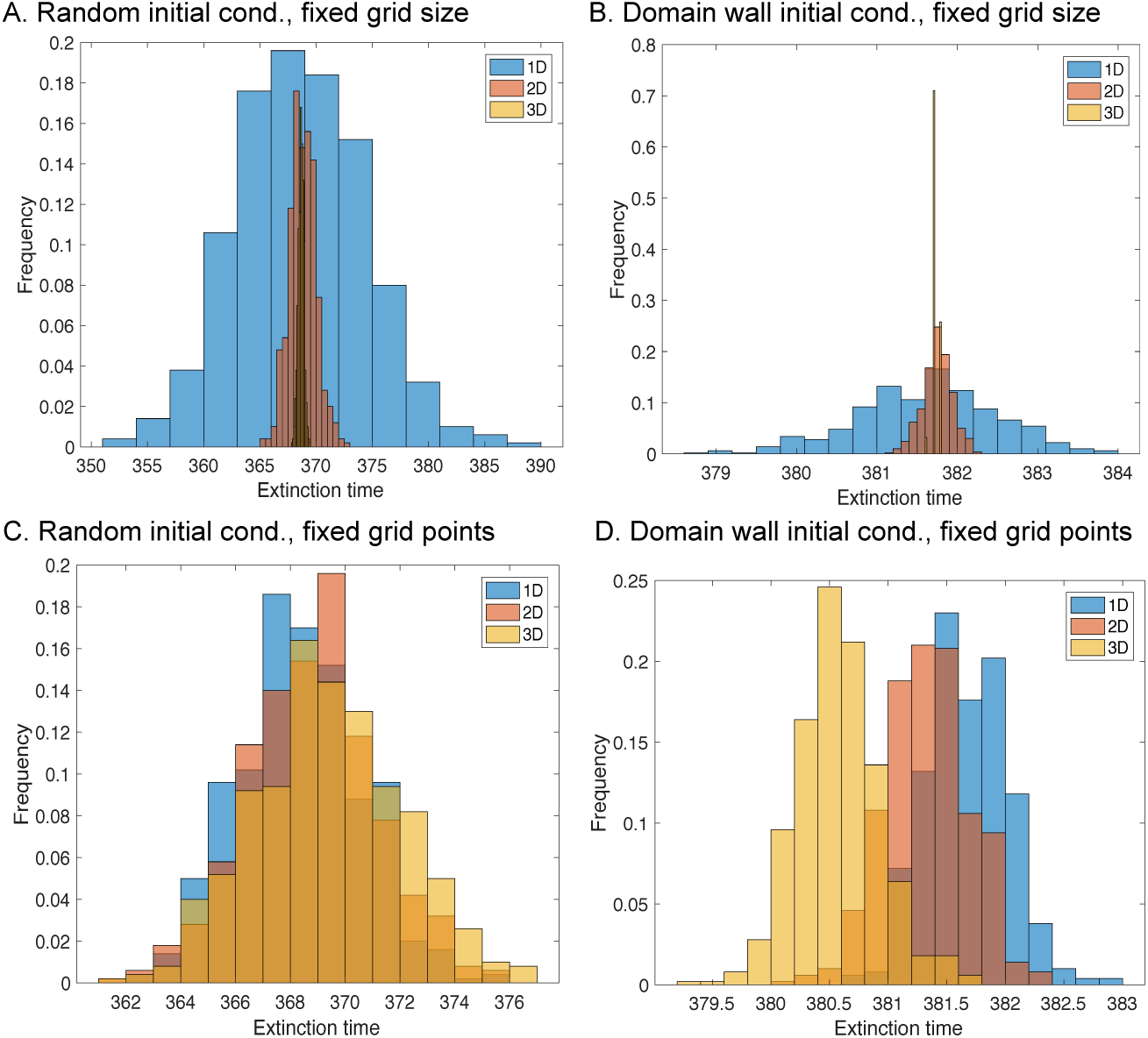
The effects of dimensionality and initial conditions. Comparison over dimensionality and different initial conditions with exit criterion *ε*_exit_ = 10^−8^. **A** Random initial condition with equal step size Δ*x* = 0.5. **B** Domain wall initial condition with equal step size Δ*x* = 0.5. **C** Distribution of extinction times by dimension with equal number of grid points. **D** Domain wall initial condition with equal number of grid points. Dimensionless parameters used γ*_U_* = γ*_V_* = 0.5, *a* = 0.9, *b* = 1, *c* = 0.5, *r* = 1, *ϵ* = 2 × 10^−3^.

### Structured initial conditions stabilize the population and extinction times increase

How are extinction times influenced by non-random initial conditions in settings of different dimensions? The impact of structured initial conditions is particularly relevant to biological processes where spatial assortment can occur in populations with limited dispersal. Therefore, we examined how the extinction or fixation times were affected by more coherent, non-random starting conditions (compare Figure 1). Figure 3 D shows the increased extinction time distribution when using a unimodal initial condition in one, two and three dimensions. A concentrated field of producers is more robust to extinction. In addition, this localized concentration has increasing density of neighbors as the dimensionality increases, which is why we see that producer extinction takes longer in higher dimensions.

Even with very structured of starting conditions–the domain wall–the deviation of mean extinction times observed (Figure 3 D) is no larger than 10%. All final states are homogeneous and correspond indeed to the stable fixed point of a non-spatial model. We thus investigated analytically the time needed to reach an equilibrium, or fixed point, using the non-spatial ODE model. To this end, we extracted an approximation which makes it possible to compare the ODE approach to the spatial PDE model. This approach allowed us to quantify the impact of spatial heterogeneity on timing to extinction.

### The predictive power of a non-spatial approach

Numerical integration of the spatial model suggested that a non-spatial analysis could be used to determine the time scale of fixation, e.g. when public good producers go extinct. This change to a simple model system should be meaningful because all final states are homogenous in space (Appendix)–the long-term dynamics of te spatially explicit system are spatially invariant. The spatially invariant version of our dynamical system is given by

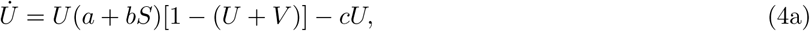

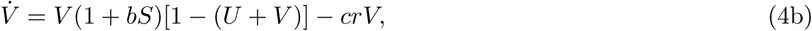

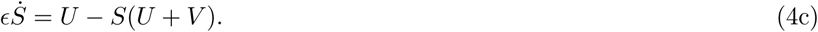

First, let us turn to the possible fixed points and their stability in the non-spatial setting. The system (4) exhibits three main steady states which exist over a wide parameter range. First, we have the mass extinction state (0, 0, *S*\*). Second, we have the case in which producers win 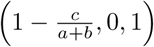. Third, we have the case in which free-rider cells win (0, 1 – *cr*, 0). Figure 4 shows examples of the dynamics between these the three main steady states in the (*U, V*) plane. Additionally, a sample trajectory is shown, which indicates the approach of a slow manifold that is be inherent to all trajectories (if this manifold exists). We can exploit this slow manifold dynamics to estimate the fixation time to the free-rider state. In addition, the linear stability conditions of the three main steady states can be calculated (see S1 Appendix for details):

- *The mass extinction state* (0, 0, *S\**) is stable if 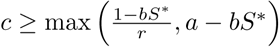.
- *The producer-only state* 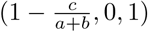 is stable provided that 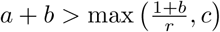.
- *The free-rider-only state* (0, 1 – *cr*, 0) is stable provided that 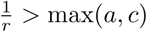.

**Figure 4:**
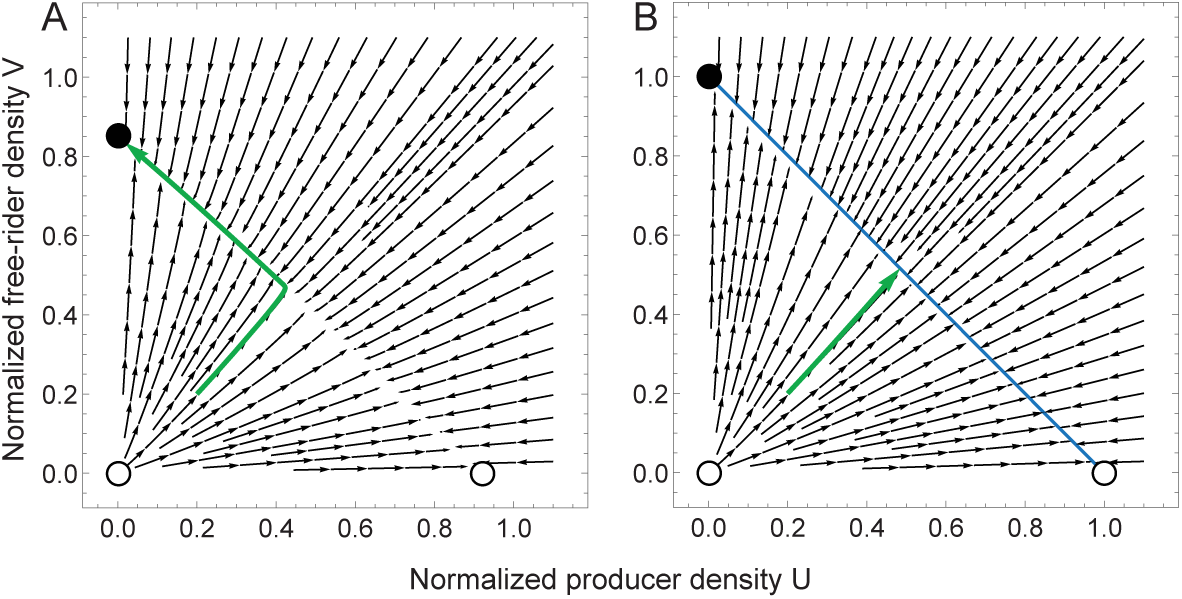
ODE phase diagram. The isolated fixed points are labelled by thick circles (stable) and hollow circles (unstable). In both cases, a curve connects the two nontrivial fixed points. **A**, phase diagram with values *a* = 0.9, *b* = 1, *c* = 0.15. The nonzero death rate *c* has caused the degeneracy of non-isolated fixed points to collapse, leaving behind a slow manifold along which the dynamics travel. **B**, phase diagram with values *a* = 0.9, *b* = 1, *c* = 0.0. The blue line is the infinite family of solutions along the curve *U* + *V* = 1.

It is interesting to note that in the case of equal death rates (*r* = 1), the producer-only state is necessarily unstable since it is assumed that production of the good comes at a cost (*a* < 1). It then follows naturally that even if we unilaterally lower the death rate of producer cell, *r* ≤ 1 the producer-only state remains unstable.

There also exist special cases for certain parameter values which involve non-isolated fixed points, illustrated by the following special scenarios. First, if *a* = 1, (*κ* = 0) and *r* = 1, (*μ_U_* = *μ_V_*), i.e. when there is no cost of public good production, an infinite family of solutions exist if 1 + *b* ≥ *c*, and is given by the condition

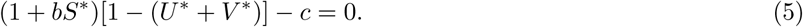

Second, if *c* = 0, (*μ* = 0), then an infinite family of solutions is given by (*U\**, 1 – *U*, U\**). Last, when the relations (1 + *b*)/(*a* + *b*) < *r* < 1/*a* and *c* < (1 – *a*)/(*r* – 1) hold, there exists a unique coexistence point of the form

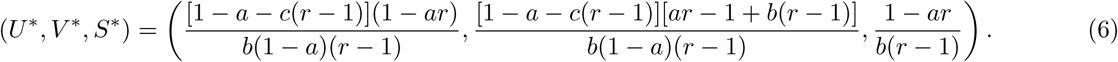

In this rare case of an unstable or saddle point equilibrium in which producer-free-rider coexist, we have *U\** > *V\** if *a* > 1 + *c*(1 – *r*). In dimensional parameters, this means 1 –*κ/α* > 1 + *μ_U_* / (1 – *μ_V_/μ_U_*). Hence, under the assumption that free-riders die faster than producers, the latter are expected to be more abundant than free-riders if the cost is smaller than the (positive) difference in their death rates, *κ* < *μ_V_* – *μ_U_*.

#### Slow manifold evolution

The systems evolution along the slow manifold is key to for the characterization of the transient dynamics of the system. Our simulations show that after a short amount of time, the average concentration approaches a curve on which it spends most of its time (Figure 2 and Figure 4A). This curve is the slow manifold. In general, this is difficult to calculate, but in certain parameter regions, we can make estimations that allow for an approximate calculation (for details see S2 Appendix).

We define the time to extinction *T_U_* and *T_V_* by the amount of time it takes for producers and free-riders to go extinct, respectively. Their non-dimensional counterparts are denoted by *τ*. In numerical procedures we specify an extinction event to occur at the threshold distance from an all-*U* or all-*V* state, given by *ε*_exit_ ≪1. Using Eq. (S2.11) with Eq. (S2.1), we obtain the estimation for the (non-dimensional) fixation time

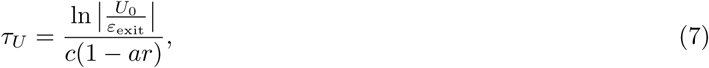

when *c* ≪ 1, that is the producer cells’ death rate is small compared to proliferation rate. In dimensional variables, the extinction time of this case can be converted easily

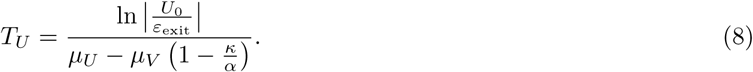

For the other cases below, the corresponding expression in dimensional variables is somewhat unwieldy. In the case when producer cells’ death and proliferation rates are of comparable magnitude, 1 – *c r* ≪ 1, we can use Eq. (S2.11) with Eq. (S2.5) to obtain the estimation for the time to producer extinction

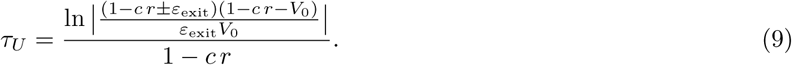

If producers win, the extinction time of free-riders (in the case of *cr* – *c*(1 + *b*)/(*a* + *b*) ≪ 1) is given by

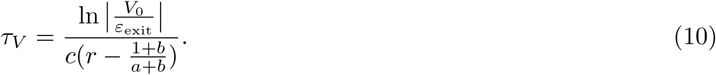

That is, the extinction time of free-riders, starting with a certain initial density of free-riders *V*_0_ grows at most logarithmically with that density. We can also see that the choice of exit criterion, *ε*_exit_ ≪ 1, which defines when the dynamics reach an arbitrarily small neighborhood of the fixed point, also grows logarithmically as ln |1/*ε*_exit_|, and not exponentially or with a power law as one might expect. As a result, one might derive some confidence in the measured *ε*_exit_-fixation time [8] as we have defined it in this paper, especially since fixation time often refers to the mean fixation time of an individual based Markov chain model of co-evolutionary dynamics [26, 27, 28, 29, 30, 31]. Next, we consider the case when *c* = 0, in which the deterministic dynamics stops on a line of non-isolated fixed points (as shown in Figure 4 B). In this case, coexistence is possible.

#### Coexistence phase

The coexistence phase can only occur when *μ_U,V_* = 0, and it is degenerate. By degenerate we mean that the state is destroyed by any small perturbation in any relevant parameter, for example a perturbation from *c* = 0 leads to the destruction of a coexistence phase, see Figure 4. However, one can still derive useful predictions for the time to coexistence (for details see S3 Appendix). Using Eq. (S3.2) with Eq. (S2.10) we obtain an estimation for the coexistence time

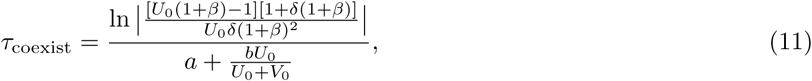

where we have defined 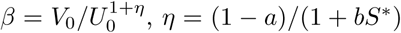 and *S\** the spatially invariant solution of the public good, governed by Eq. (4c).

#### Random spatial heterogeneity has little impact on extinction times

Our first-order approximations for the extinction time provide useful insight into the sensitivity of extinction times on the parameters. For example, the extinction time is inversely proportional to death rate and production cost, but directly proportional to the birth rate. Also, it is only logarithmically dependent on the initial concentrations and the exit threshold. The influences of *b* = *ρ/δ* (the ratio of public good production over public good consumption rates) and *ϵ* = *α/δ* (the ratio of proliferation rate over public good consumption rate) entered the approximation at higher orders and are therefore subdominant. While typically *b* ≈ 1, *ϵ* can often be assumed to be a small parameter–within a typical cell cycle the public good produced is also quickly consumed. Overall, this slow manifold approximation can be used to predict extinction and coexistence times very accurately, see Figure 5.

**Figure 5:**
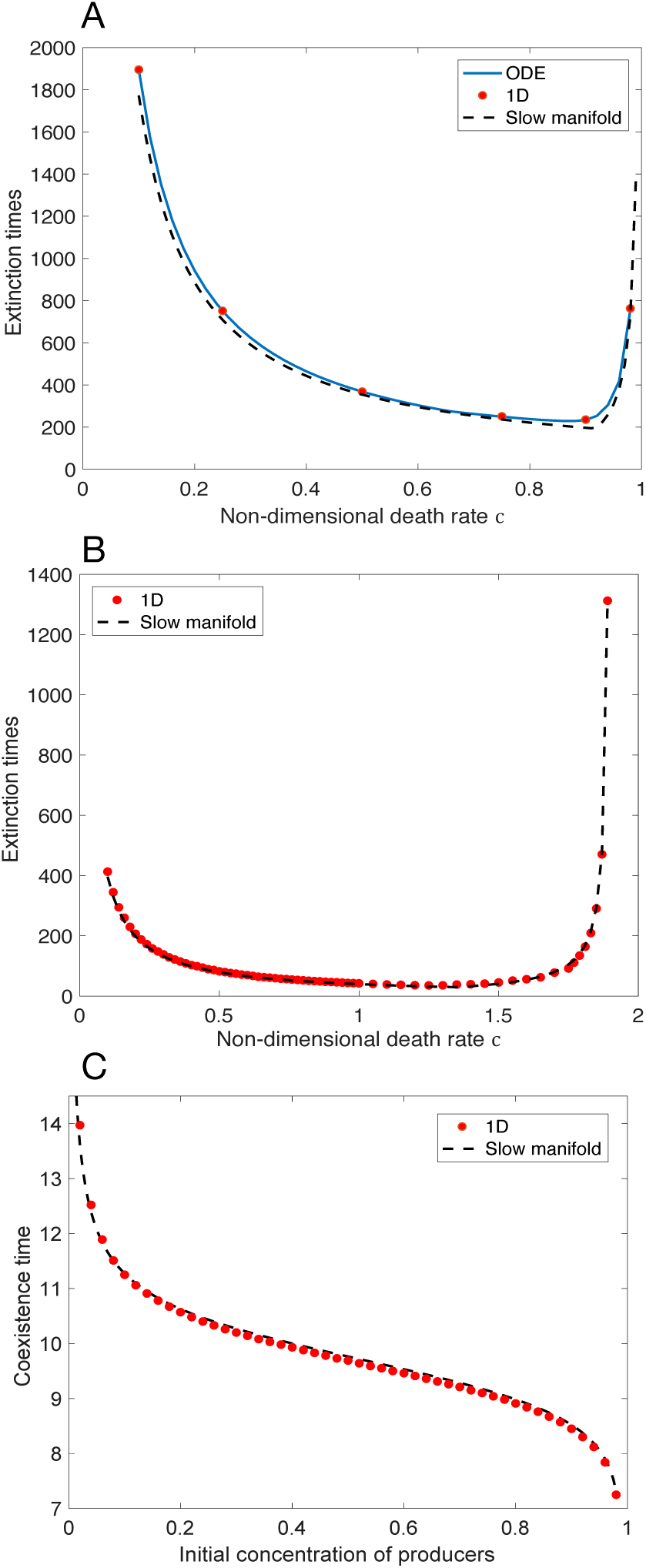
Extinction and coexistence times using random initial conditions. **A** The time to within a *ε*_exit_-radius of the stable fixed point as a function of the non-dimensional death rate of producers *c* = *μ_U_ / α*. Other non-dimensional parameters used: *a* = 0.9, *b* = 1, *r* = 1, *ϵ* = 2 × 10^−3^. **B** The time to within a *ε*_exit_-radius of the producer-only state, as a function of *c*, with *r* = 1.2 and all other parameters as before. **C** The time to within a *ε*_exit_-radius of the coexistence manifold as a function of the initial concentration of producers (the initial concentration of free-riders is *V*_0_ = 0.01). The 1D spatial is taken using random initial conditions averaged over 100 simulations (the standard deviation of times is less than the point size). Parameters used: *a* = 0.9, *b* = 1, *c* = 0.5, *ϵ* = 2 × 10^−3^. In all panels, the spatial system was set up using random initial conditions and averaging over 100 simulations (the standard deviation of fixation times was less than the point size chosen in the graphs). Due to the re-scaling of the non-dimensional system, all times can be understood in units of the average cell cycle length (1/*α*).

In Figures 5 A, B, we compare the extinction times calculated with the dimensionless framework of system (4), to simulations of the full system (1D), and to the slow manifold approximation, Eq. (9), in cases of producer extinction. For the case of coexistence, Figure 5 C, *c* = 0 (or *μ_U_* = *μ_V_* == 0), our approximation (11) is in very good agreement with the numerical simulations. Remarkably, this time to coexistence is orders of magnitude smaller than extinction times, and relatively robust to changes in the initial concentration of producer cells, if we keep this concentration above 20 and below 80%. For much lower or much higher initial producer concentrations, this time rapidly increases or declines, respectively.

### On a structured domain, the domain length can have a strong impact

To investigate the impact of the domain length, we considered uni-modal, domain wall, and random initial conditions for the concentration of producer cells, free-rider cells, and the public good. Simulations showed that the domain length, when given purely random starting conditions, did not impact the extinction time. However, for the domain-wall and other structured conditions, the size of the domain influenced the fixation time. The invasion of free-riders into the space occupied by producers is similar to traveling waves observed in standard Fisher equations [32, 33]. From this, we hypothesize that the total extinction time is modified by the time it takes for this wavefront to reach the end of the unstable region. We can thus say that the extinction time can be represented by the following linear combination

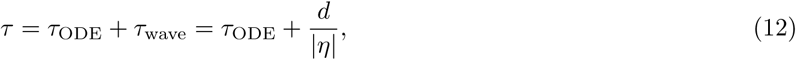

where *d* is the distance travelled by the Fisher wave, and |*η*| the speed of the wavefront.

#### free-rider invasion

In the case of an unstable producer-only state, a free-rider population initially separated will invade the producers. We consider a domain where a boundary exists between free-riders and producers. Simulations show a traveling wavefront of free-riders into the producer-only region. An approximation to the speed of the wavefront is given by (see S4 Appendix for details)

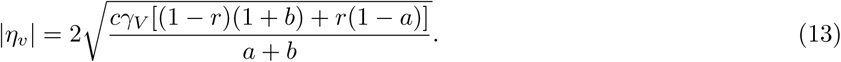

Hence, the total time to extinction for *c* ≪ 1 is given by

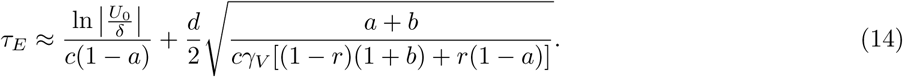

Note that this approximation is valid for *c* ≪ 1. In the case of *c* ≈ 1 we would replace the first term of the approximation (14) by the right hand side of Eq. (9). We tested this prediction against different parameter values (see Figure 6 A) and found good agreement with the prediction. The time added to the ODE prediction can be described by the amount of time needed for the free-rider wavefront to travel the distance necessary to cover the entire finite domain.

**Figure 6:**
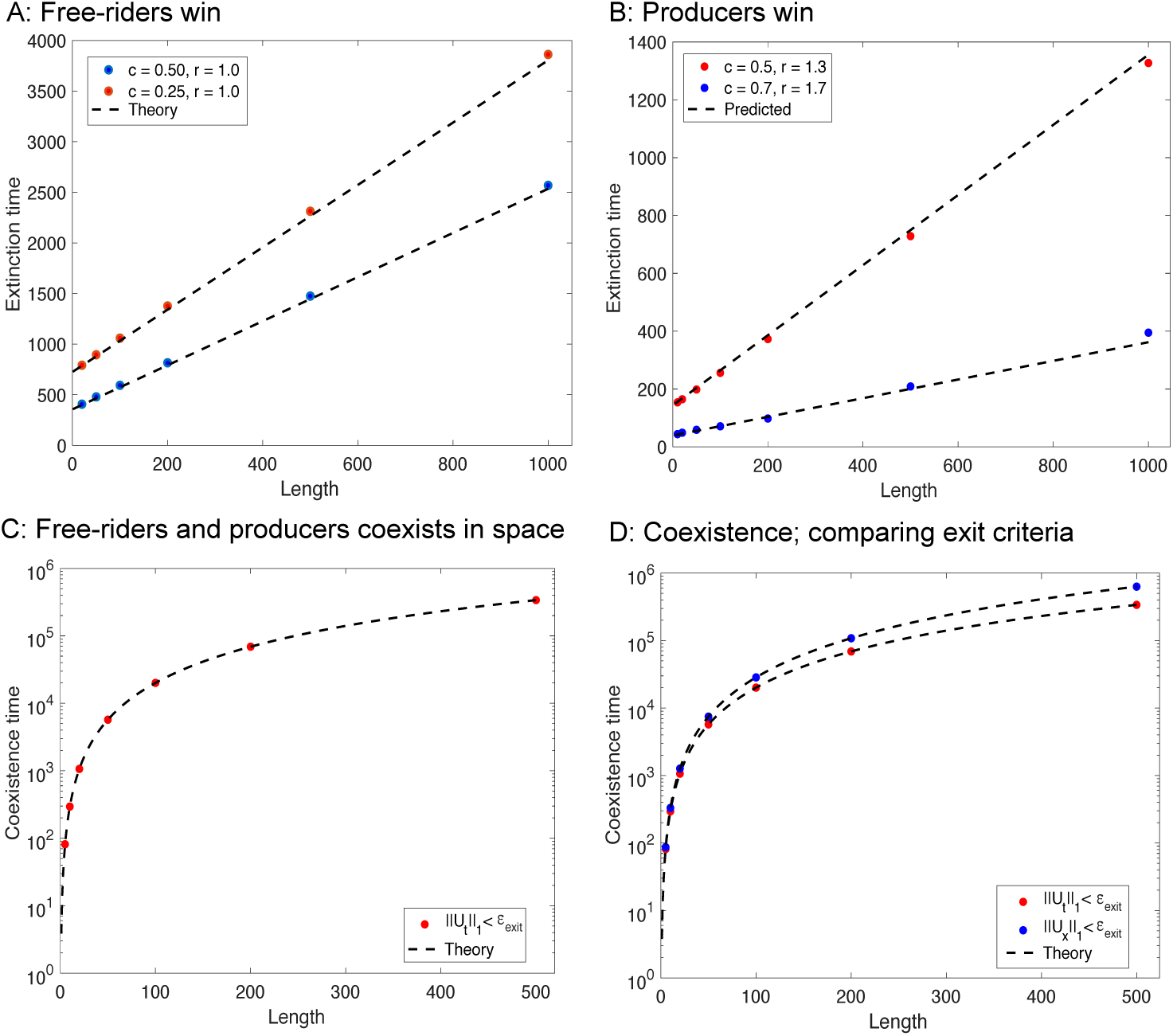
Extinction times increase linearly with length, and coexistence times increase logarithmically with length. **A**: Producer extinction time vs. length of the domain with *r* = 1. theoretical lines were obtained using Eq. (14). **B**: free-rider extinction time vs. length of the domain. theoretical lines were obtained using Eq. (16). **C**: Coexistence time vs. length with *c* = 0, with theory obtained from Eq. (S5.3). **D**: Comparison of exit criterion on the calculated time to coexistence. The system was considered to have reached a fixed point when the producer density was within the *ε*_exit_-neighborhood of that fixed point. We compare the exit criteria for the temporal (S5.3) *p*-norm (red) with the spatial (S5.4) *p*-norm (blue) with, using *p* = 1. Other (non-dimensional) parameters used in all panels: *a* = 0.9, *b* = 1, *γ_U_* = γ*_V_* = 0.5, *ϵ*= 2 × 10^−3^. Note that we here show the dimensionless domain length, which scales as the square rood of public good diffusion constant over public good consumption rate, Γ*_G_/δ*.

#### Producers invasion

In the case of an unstable free-rider-only state, a spatially separated producer population will invade the free-riders. Unlike the free-rider invasion, the producer invasion is more complicated. This is mostly due to the impact of *r* on the location of the free-rider-only state. Suppose we are in a region in parameter space where the free-rider-only state is unstable. If this state is in the region (0,1), we can proceed as we did in the previous section. However if 1 – *c r* < 0, which is biologically feasible, the wavefront travels from the mass extinction state, rather than the free-rider-only state as before. This is reflected in the different wave speeds obtained below. Thus, an approximation to the speed of the wavefront is given by (see S4 Appendix for details

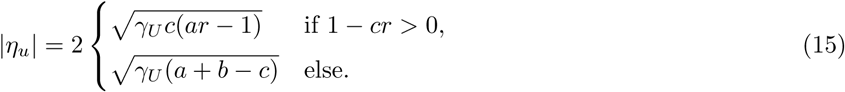

We note that the first case is valid only if 1 – *cr* > 0. This is due to the free-rider state moving into the negative region and hence is nonphysical. The appropriate unstable point is then the extinction state and the speed is given instead by the second case. Hence, the total time to extinction for *c* ≪1 is given by

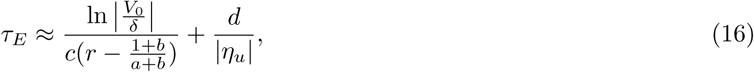

We tested this prediction against simulations for different parameter values and found good agreement with Equation (16), see Figure 6 B.

#### Diffusion times are relevant only in the absence of cell death

The special case of vanishing cellular death rates *c* = 0 and *r* = 1 revealed the possibility of coexistence of producers and free-riders along a one-dimensional subset of the state space. In this context, it is interesting to examine the limit as *c* → 0 in an initially highly structured population where producer and free-rider cells are highly segregated at time 0. In this case, the wavefront speed, e.g. of an expanding free-rider population, tends to 0. A vanishingly slow wavefront would imply that the extinction time tends to infinity, *τ* → *∞*, which seems unrealistic. Indeed, the diffusion time that governs cellular dispersion becomes relevant. Scaling implies that the characteristic time for diffusion is *τ*_diffusion_ ~ *L*^2^/*D*, where *D* is the diffusion coefficient of both types of cell. For *c* > 0, the wavefront should move faster than diffusion, and so this type of scaling with the diffusion constant is not seen for finite cell death rates. Cellular diffusion should be the driving factor in long times as *c* → 0. To test this, we considered *c* = 0 and calculated the time to homogeneity. An analytical expression was calculated in S5 Appendix. The comparison between domain length and the time scale to reach diffusion driven coexistence in this special case of diffusion dominated extinction is shown in Figures 6 C and D. The time to coexistence increases rapidly with small domain length, and less so when the spatial domain is large.

## Discussion and Summary

Here we considered a spatial linear public goods population game model in its deterministic form. We have investigated the impact of spatial arrangement of public good producer cells and free-rider cells on the temporal scales of extinction or coexistence during the co-evolution of these populations on a finite spatial domain. The model typically exhibits fixation for either producers of the public good or free-riders, which critically depends on the frequency-dependent birth and death rates, and on properties of the public good itself, such as the cost of production. While the cost to benefit ratio plays a part in this, the overall dependencies can be more involved. Our analysis has shown that the type of initial condition, in terms of the initial population placement in space, has a large impact on the predicted time to fixation. The dynamics of unstructured (random) initial conditions can be captured by an non-spatial ODE-model, for which extinction times can be calculated analytically. For structured initial condition, e.g. a domain wall, the taking over of one cell type on a finite spatial domain increases the extinction time linearly with the size of the spatial domain on which the population can grow.

Our numerical simulations show that all spatial inhomogeneities are ultimately removed, but not insignificant in regards to the time it takes to reach spatial homogeneity. We have to this point, stated without cause that the time to the slow manifold is subdominant with respect to the slow manifold, and in 1D, the wavefront time. The coexistence time calculation when cellular death is neglected provide an insight into this phenomenon. Consider the difference in time scales from figure 4, the extinction time is much larger in magnitude compared to the coexistence time. Since the coexistence points exist as a line connecting the two nonzero fixed points, the coexistence time can also be thought of as the time it takes to reach the slow manifold. To understand this better, consider 0 < *μ* ≪ 1. The line of fixed points has been removed leaving the boundary fixed points (and possibly an unique coexistence point). The collapsed line of fixed points retains some of its impact on the time and we recognize the line as an approximation to the slow manifold. Hence the coexistence time is an approximation for the amount of time the trajectories spend before reaching the slow manifold. The orders of magnitude between the two time scales explains why we can neglect the time to reach the manifold.

Our results highlight a point often ignored in the evolutionary dynamics literature, which typically focuses on the evolutionary stable states (ESS) and ignores the temporal dynamics of selection. Similar tendencies are apparent in the wider field of the study of ecological systems, where transient behavior has often been secondary to determining long-rem stable states [34]. Our analysis shows that both population dynamical parameters, such as death rate, the initial condition, and the spatial extent of the population influence the time it takes to reach the ESS. These results are particularly relevant to cancer, where public goods might be a common feature of tumor-ecological stability, for example as seen by the evolution of autocrine growth factor production [11].

The tumor public goods game investigated in Archetti et al. [35] uses the growth factor IGF-II as the public good. One can parametrize our system to describe that experimental system in the following way:

- The diffusion coefficient of IGF-I (a protein very similar to IGF-II) is given by Γ*_G_* = 1.21 × 10^−6^ cm^2^/s [36].
- Diffusion of cells is typically assumed to be on the order of Γ*_U_* =Γ*_V_* = 10^−8^ cm^2^/s [37].
- If we assume a base line doubling time of the cells to be 24 hours we get a growth rate of *α* = 8 × 10^−6^ s^−1^.
- While cancer cells often have more moderate net growth rates (*α/μ* ≈ 1), cell cultures *in vitro* have death rates around one-tenth the size of the growth rate. Therefore we assume equal death rates, *μ* = 8 × 10^−7^*s*^−1^.
- The cost can be estimated from the data in [35], by comparing the growth rate of producers vs. non-producers at high IGF-II concentrations (when the benefit is absent). This yields an estimate of *κ* ≈ *α*/4 = 2 × 10^−6^*s*^−1^.
- It is known that the half-life of IGF-II in tissue is on the order of 10 minutes, yielding a rate *δ*= 10^−3^ ng/cm^3^ s^−1^.
- The production rate of IGF-II can be estimated from experimental data to be *ρ* = 10^−3^ ng/cm^3^ s^−1^ [38].

Using these values in our model leads to dimensionless parameters

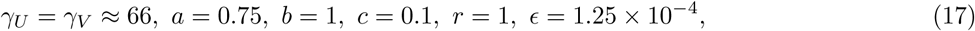

which are comparable with most of the parameter values used in this manuscript. Exceptions are *γ_U_, γ_V_* whichare around 100 times higher in magnitude in this example. However, our simulations with initial conditions of random cell placement showed no real differences, except for initially highly structured populations. In the most extreme case of initial segregation by cell type, the parameters *γ_U_, γ_V_* determine the speed of the wavefront, e.g. of a free-rider population that takes over, which influences the time to extinction of producer cells. The qualitative behavior of the transient dynamics, however, remain the same.

In practice, it can be difficult to calculate the biological parameters that are relevant to public good-driven evolutionary dynamics, such as the consumption and production rates of the public good, or its context dependent diffusion parameters. Other important rates, such as cellular death and birth rates are perhaps easier to infer, and the cost of production can be quantified in relation to the average producer cell’s proliferation rate [39, 35]. Based on our insights here, we propose a novel experiment which exploits our analytical solutions. The goal is to estimate the biological parameters of cellular public good games.

Suppose that one is interested in estimating all dimensional parameters. There there are nine such parameters (Table 1), which would require nine equations or conditions. According to the results of this manuscript, these conditions can be realized using varying initial cellular configurations in space. One can first place only producers onto a rectangular plate and wait until the population has reached the carrying capacity on the entire domain, recording the population size and the level of public good produced (here after labeled as growth factor). The same can be be obtained for free-riders to determine their final size. We would then propose to partition the rectangular plate into two parts and to place an initial concentration of producers and free-riders *U*_0_, *V*_0_, respectively, into separate partitions. We deine *t* = 0 as the time when the partition is removed, and record the time *t*_1_ = *T_f_* that one subpopulation wins (the other has gone extinct). Repeating this experiment with eight more different sized partitions will lead to measuring subsequent extinction times *t*_2_,…, *t*_9_. Supposing that free-riders win throughout these different experimental conditions, using Eq. (14), we have

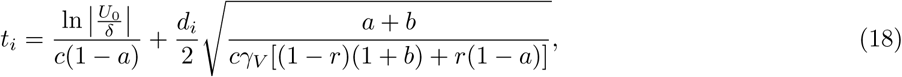

for the different conditions *i* = 1, …, 9. This formula can be converted to dimensional variables. Using any routine numerical solver, the parameters to fulfill these equations, and their likelihoods, can be estimated. By using multiple runs with the same partition sizes, one can get more accurate measurements and be more confident in the obtained values. Hence, an estimation of the metabolic cost associated with the production of the good can now be calculated. A similar process can be carried out if producers win.

In Summary, we have considered the spatial growth dynamics of producer and free-riders, determined by a diffusible public good, in one, two and three dimensions. Extracting a slow manifold solution, we obtained a good estimate for the time to extinction of a cell type. For invading populations, i.e. for initially highly segregated sub-populations, we observed a traveling wave solution. We thus calculated the estimate of the wavefront speed and added the corresponding time to the previously calculated extinction time, which was in good agreement with numerical simulations. Our spatial model can be used to generalize the tumor ecological dynamics presented in [17], which was used to assess adaptive anti-cancer strategies assuming a well-mixed population. Our spatial considerations can help refine such models and provide more accurate predictions, which could reveal critical new information with regard to the time scales of population transformations.

## Acknowledgments

We thank Robert S. Gatenby for fruitful discussions and useful comments.

## Supporting information

### S1 Appendix. Linear stability analysis

We analyze each steady state in turn, but first we need the Jacobian

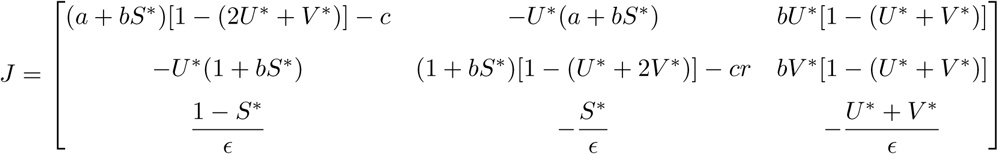

The mass extinction state (0, 0, *S*\*):

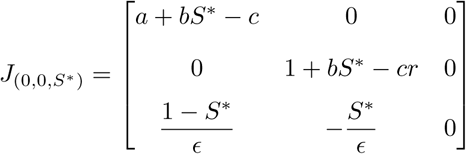

The eigenvalues can be read off λ= 0, 1 + *bS* – cr, a* + *bS* – c*. The state is stable if 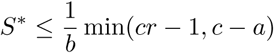.

If *U* wins the game:

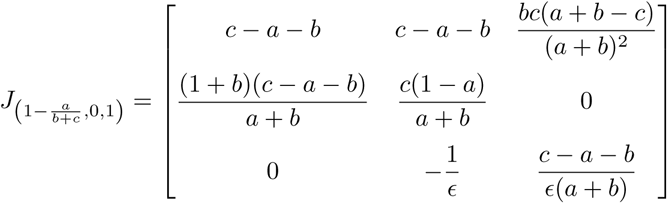

The eigenvalues are 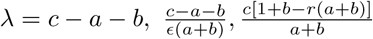. The state is stable if 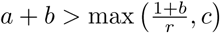 (note that this requires *r* > 1 since we assume *a* < 1).

If *V* wins the game:

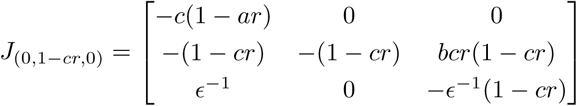

The eigenvalues are λ = –*c*(1 – *ar*), –(1 – *cr*), –(1 – *cr*)/*ϵ* and so this state is stable if 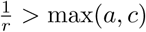.

A special case is given by *c* = 0 with *U*\* + *V*\* = 1 and *S*\* = *U*\*. The Jacobian simplifies considerably,

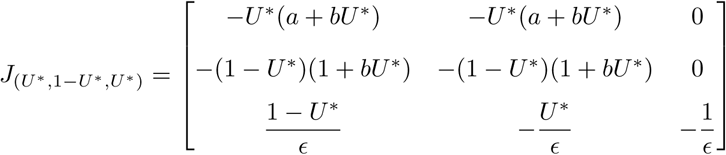

The eigenvalues are all non-positive, 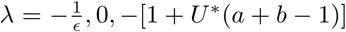. Hence the coexistence state when *c* = 0 is stable.

### S2 Appendix. Extinction/Fixation times calculation

A slow manifold reduction can be made in parameter regions where one eigenvalue is sufficiently smaller than the other. This was done for *c* ≪ 1 and 1 – *cr* ≪ 1. We suppose we may neglect the dynamics of the growth factor since *ϵ* ≪ 1, we set *ϵ* = 0. The eigenvector is 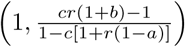. Numerical simulations indicate that the linear terms are sufficient, but we include the next order as well.

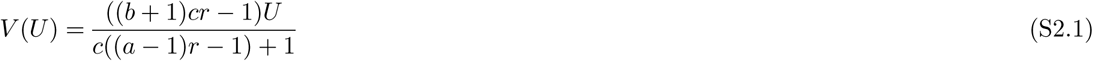

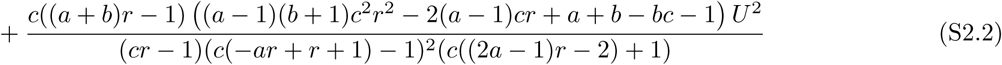

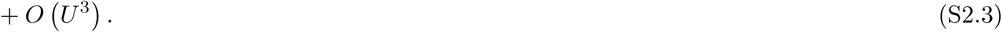

Inserting this into the equation for *U* we have the dynamics on the manifold for *U* ≪1 given by

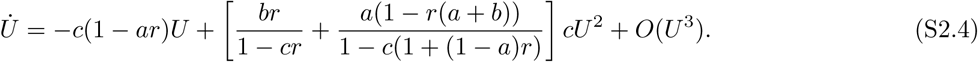

If instead we are in the other regime with 1 – *c* ≪ 1, the eigenvector is (0,1), we obtain

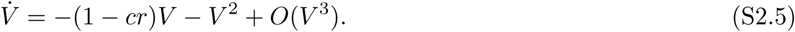

When producers win, we can play the same game to obtain an estimation for the slow manifold and the corresponding time. When *c*(1 + *b*) ≪ (1 + *cr*)(*a* + *b*), we can estimate the slow manifold

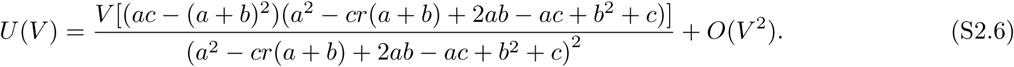

For *V* ≪ 1, the dynamics then evolve according to

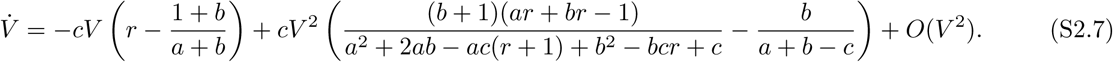

When *c* ≈ *a* + *b*, (the other eigenvalue is small), we obtain at leading order

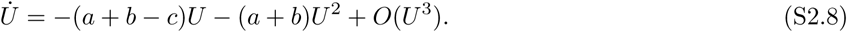

Let us now remove the restriction that *ϵ* = 0 and consider the general solution and see the effects of nonzero *ϵ* on the fixation time. With the aid of Mathematica, we obtain the slow manifold for small *ϵ* with *c* ≪ 1

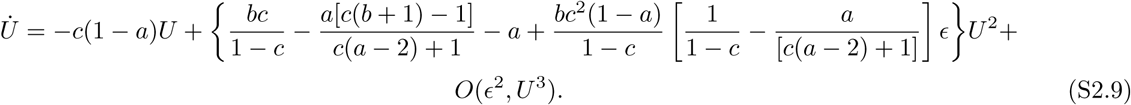

Therefore, the effects of the growth medium equation are only small corrections, on the order of *ϵ* as they only enter into the *U*^2^ term. A similar calculation can be done when *c* ≈ 1.

Linearization near fixed points give eigenvalues and eigenvectors that show how trajectories will approach the point (if it is stable) and how long it will take. Sometimes, it is possible that one eigenvalue is much smaller than the other. When this happens all trajectories heading towards the fixed point collapse onto what is known as the slow manifold. In this case, the dynamics can be reduced to a single dimension and we can discern general cases where the time to fixation can be calculated analytically. Assuming a slow manifold expression exists for *V* as an expansion in *U*, a general approximation to the dynamics is given by

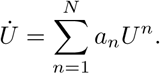

The easiest and most common are those where the linear manifold or next order is sufficient. We do the *N* = 2 case the *N* = 1 follows as a consequence, the higher order ones are more complicated, but doable if necessary. We assume the initial value *U*_0_ and we can approach the fixed point *U*\* exit criterion radius *ϵ*_exit_ from either side

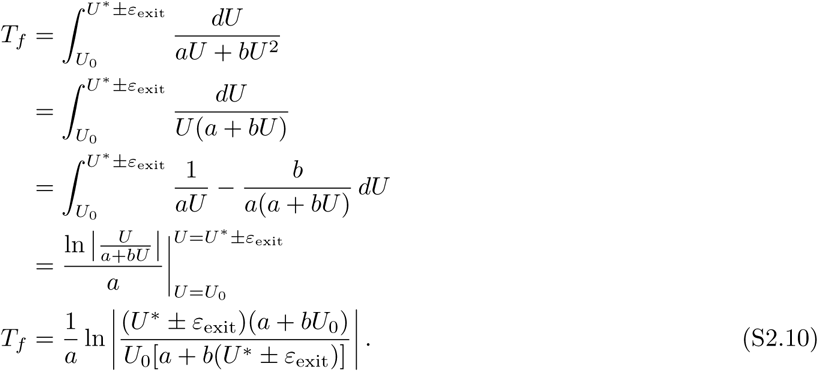

If *b* = 0 we have

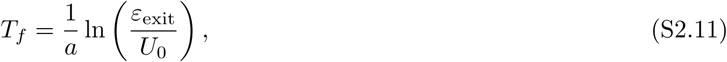

while if *a* = 0 L’Hopital’s rule gives

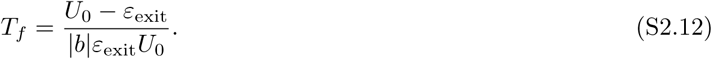

### S3 Appendix. Coexistence times calculation

With *c* = 0 we can divide Eq. (4a) by Eq. (4b) and arrive at

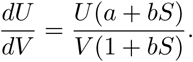

If we make the assumption that *S* equilibrates rapidly, we can substitute *S*\* = *U*_0_/(*U*_0_ + *V*_0_). This allows us to solve this equation for *V* and arrive at

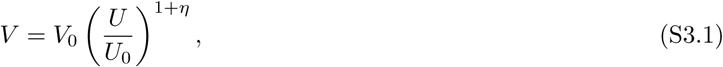

with 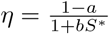. Inserting (S3.1) into (4a), we have

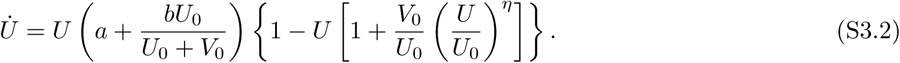

We also make the observation that with the accepted parameter values for *a* and *b* we see that *η* ≤ 0.1 and so it is a small parameter. For *t* → ∞, a steady state is given by the solution to

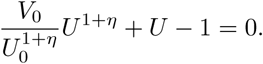

Expanding *U* in *η* ≪ 1 we obtain

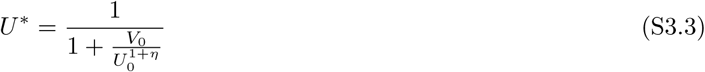

### S4 Appendix. Calculation of wavefront speed in 1D and the corresponding extinction time adjustment

Though no traveling wave solutions can exist in the long term, nor were spatially inhomogeneous solutions observed, we can still calculate the time it takes for an population of free-riders to invade from one side of the domain to the other.

Following the work in [33], we seek an approximation to the speed of the wavefront. The instability of the unstable state drives the dynamics. To this end we consider the linearization about that unstable steady state given first by 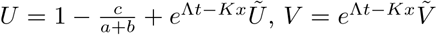 and 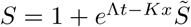. Plugging this into Eq. (3) and neglecting higher order terms, we arrive at a linear system **A**(Λ, *K*)**p** = 0 where 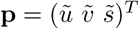. We are interested in nontrivial solutions to this linear equation, which implies we are looking for when the determinant is 0. This leads to three separate values for Λ

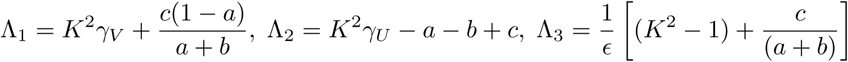

The only relevant value is the first one. Since we are interested in the speed of the front, we are looking at

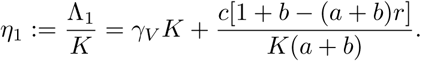

Minimizing over *K*, we arrive at

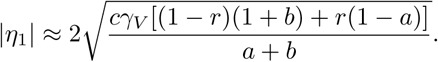

If instead, we linearize about the consumer-only state, we use 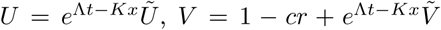 and 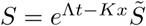. In an analogous manner to the producer state, we arrive at

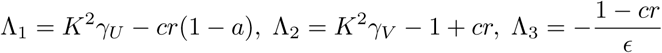

It again can be shown that only one of these values leads to instability and so the speed

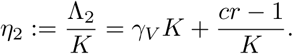

Minimizing over *K*, we arrive at

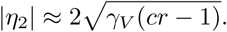

Let *d* be the size of the domain of producers, then the wave’s travel time is given by 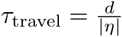. In the case of an even split of producers and free-riders *d* = *L*/2 and we arrive at

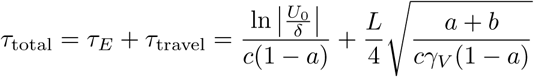

In the case from the text, we have *γ* = 0.5, *a* = 0.9, *c* = 0.5, we arrive at

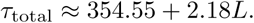

### S5 Appendix. Calculation of coexistence time

In the case *c* = 0 (no death), the ODE model gives an approximate time for coexistence to the line *U* + *V* = 1. Simulations of the spatial model show a similar rapid approach to the line with the *average* population. However, the system continues to run till it is homogeneous. Since there is no traveling wave in this scenario, the only method to reach homogeneity is that of diffusion. In a similar way to the wavefront problem, we assume that the total time is given by

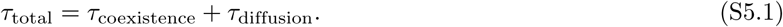

The approach of the average population to the line is rapid (Figure 5). Therefore, we make the approximation that *U* + *V* = 1 for calculating the diffusion time. This uncouples *U* and *V* and reduces Eq. (3) to

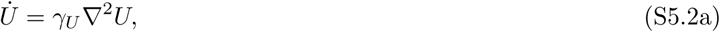

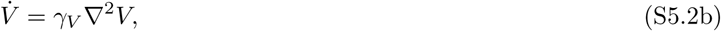

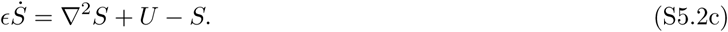

We expand the solutions of *U, V, S* using a cosine series 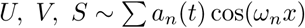, where 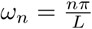. The solution for *U* is given by

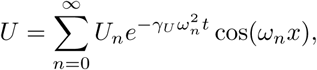

where

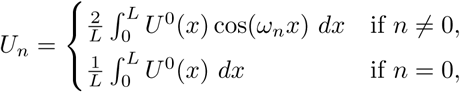

and *U*^0^(*x*) := *U*(*x*, 0). The solution for *V* is analogous and we note that the time for *S* to equilibrate is dependent on *U* and the time is exponentially small.

As an example, we choose the domain wall initial conditions *U* = *I_u_H*(*L/*2 – *x*) and *V* = *I_v_H*(*x* – *L*/2). We obtain

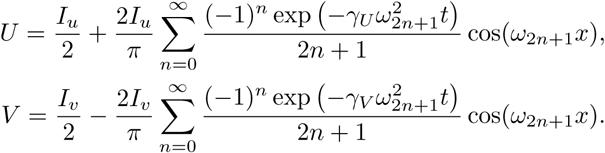

The exit criterion is satisfied when ∥*U_t_*∥_1_ < *ϵ*_exit_ and ∥*V_t_*∥_1_ < *ϵ*_exit_. Noting that *n* = 1 governs the time to homogeneity (*n* > 1 is subdominant), we obtain

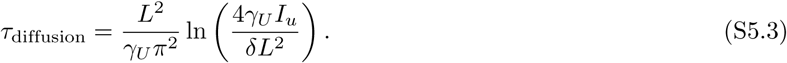

A similar result is obtained in higher dimensions. The procedure is the same for any initial condition. However, we note that the choice of the exit criterion will change the form of *τ*_diffusion_. Examples include ∥*U_t_*∥_∞_ or ∥*U_x_*∥*_p_* (the p-norm). For example, using ∥*U_x_*∥_1_ we obtain

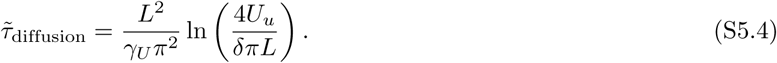

It is easy to see that this choice leads to a larger increase for increasing length. A comparison is shown in Figure 6.

### S6 Appendix. Ruling out pattern formations

We have solved Eq. (3) with different initial conditions in one-,two- and three-dimensional systems. Though the time to extinction or coexistence can vary, all systems eventually reached spatial homogeneity. A main cause of spatial pattern formations in diffusive systems is through diffusion-driven instability (DDI) which occurs through a mechanism known as a Turing bifurcation. We show here that the model does not admit a DDI via a Turing bifurcation.

### S7 Appendix. Different initial conditions converge to the homogeneous solution

Our simulations show that systems with higher initial spatial order take longer to converge to the homogeneous solution, see Figure 7.

**Figure 7:**
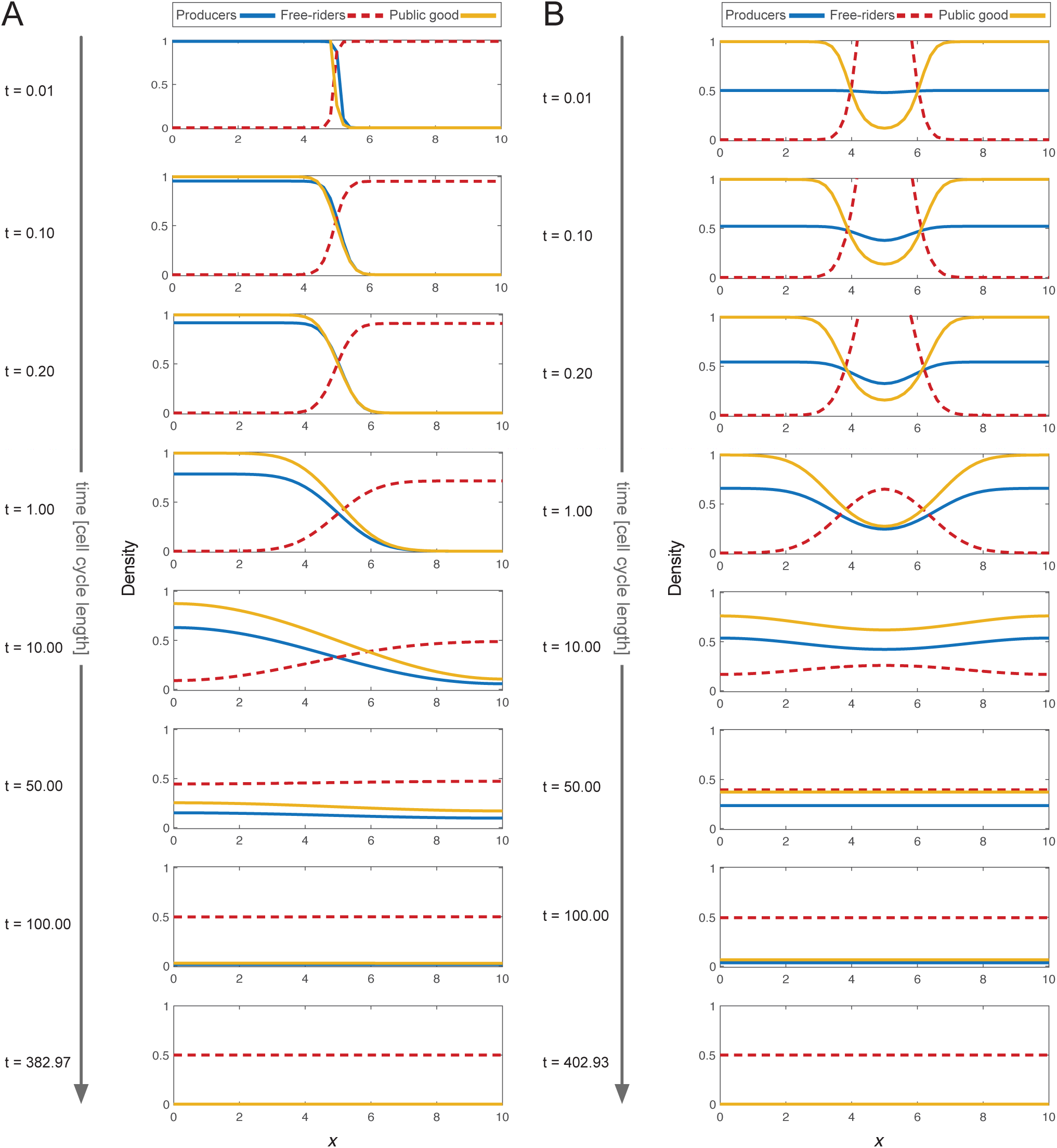
More spatially ordered systems take longer to converge to the homogeneous solution. **A**: The domain wall initial condition (Figure 2 B). **B**: Uniform producer density with a Gaussian distribution for free-riders in the center of the domain (Figure 2 C). Other (non-dimensional) parameters used in all panels: *a* = 0.9, *b* = 1, *r* = 1, *γ_U_* = γ*_V_* = 0.5, *ϵ* = 2 × 10^−3^.

